# Glycosylation flux analysis reveals dynamic changes of intracellular glycosylation flux distribution in Chinese hamster ovary fed-batch cultures

**DOI:** 10.1101/121517

**Authors:** Sandro Hutter, Thomas K. Villiger, David Brühlmann, Matthieu Stettler, Hervé Broly, Miroslav Soos, Rudiyanto Gunawan

**Affiliations:** Institute for Chemical and Bioengineering, Department of Chemistry and Applied Biosciences, ETH Zurich, 8093 Zurich, Switzerland; Swiss Institute of Bioinformatics, 1015 Lausanne, Switzerland; Biotech Process Sciences, Merck Biopharma, 1804 Corsier-sur-Vevey, Switzerland; Department of Chemical Engineering, University of Chemistry and Technology, 166 28 Prague, Czech Republic

**Keywords:** N-linked glycosylation, flux analysis, CHO cells, constraint-based modeling, monoclonal antibody

## Abstract

N-linked glycosylation of proteins has both functional and structural significance. Importantly, the glycan structure of a therapeutic protein influences its efficacy, pharmacokinetics, pharmacodynamics and immunogenicity. In this work, we developed glycosylation flux analysis (GFA) for predicting intracellular production and consumption rates (fluxes) of glycoforms, and applied this method to CHO fed-batch monoclonal antibody (mAb) production using two different media compositions, with and without additional manganese feeding. The GFA is based on a constraint-based modelling of the glycosylation network, employing a pseudo steady state assumption. While the glycosylation fluxes in the network are balanced at each time point, the GFA allows the fluxes to vary with time by way of two scaling factors: (1) an enzyme-specific factor that captures the temporal changes among glycosylation reactions catalyzed by the same enzyme, and (2) the cell specific productivity factor that accounts for the dynamic changes in the mAb production rate. The GFA of the CHO fed-batch cultivations showed that regardless of the media composition, the fluxes of galactosylation decreased with the cultivation time in comparison to the other glycosylation reactions. Furthermore, the GFA showed that the addition of Mn, a cofactor of galactosyltransferase, has the effect of increasing the galactosylation fluxes but only during the beginning of the cultivation period. The results thus demonstrated the power of the GFA in delineating the dynamic alterations of the glycosylation fluxes by local (enzyme-specific) and global (cell specific productivity) factors.

## 1. Introduction

The introduction of Quality by Design (QbD) and Process Analytical Technology (PAT) initiatives by the US Food and Drug Administration has driven a resurgence in research efforts and innovation in biopharmaceutical process development and manufacturing (FDA, 2004; Rathore, 2009). QbD and PAT prescribe continuous process improvements through well-defined process objectives, science-based product and process understanding, timely (online) process measurements, and process system design, analysis and control. In the current competitive and burgeoning market for therapeutic proteins and biosimilars, the biopharmaceutical industry faces numerous challenges in ensuring product quality, shortening time to market, improving cost effectiveness and creating manufacturing flexibility (Li et al., 2010). One of the most important quality attributes of recombinant therapeutic proteins is the structure of the asparagine-linked oligosaccharide sugar or N-linked glycan attached to the proteins. This glycan structure can influence protein folding (Aebi, 2013), secretion (Zhang and Kaufman, 2006) and stability (Arosio et al., 2013), and has an impact on bio-activity (Umaña et al., 1999), efficacy (Goh et al., 2014) and immunogenicity (Harding et al., 2010). Consequently, the regulation and control of N-linked glycosylation has received much attention from the biopharmaceutical industry.

The N-linked glycan of a protein is processed post-translationally in the rough endoplasmic reticulum (ER) and Golgi cisternae. There exists a natural heterogeneity in the outcome of the glycosylation process. The biosynthetic pathways of N-linked glycosylation processing in the Golgi involve a relatively small number of enzymes, and therefore, minor alterations in the activities of these enzymes can cause significant changes in the resulting glycan profile (i.e. the distribution of the glycan structures) of a protein, sometimes with detrimental consequences (Aebi and Hennet, 2001). The glycan profile has been shown to depend on, among other things, cell culture parameters (e.g., dissolved oxygen, pH, temperature, osmolality), culture media composition, and expression hosts (Hossler et al., 2009; Ivarsson et al., 2014). Yet, the mechanisms of these dependencies still remain unclear. Transcriptomics, proteomics and metabolomics analysis have been used to get a better picture of the intracellular changes in protein glycosylation, caused by modifications in the cell culture conditions (Sha et al., 2016). Furthermore, mathematical models of the glycosylation network have been created to explain the observed glycoform profiles and to predict the glycosylation outcome (Jimenez del Val et al., 2011; Liu et al., 2013; St Amand et al., 2014; Umaña and Bailey, 1997) Recent modelling efforts have focused on linking extracellular metabolite measurements to the protein glycosylation process by coupling the dynamic model of the bioreactor with the intracellular model of glycan metabolism and glycosylation network (Jedrzejewski et al., 2014). However, these detailed models comprised partial differential equations and possessed a large number of unknown kinetic parameters, posing a challenge in model simulations and parameter estimation.

On the opposite spectrum of modelling framework, the constraint-based modelling (CBM) uses a simple and parameter-free approach, requiring only the stoichiometry of the reactions to make predictions on the reaction fluxes. The most successful application of CBM has been in the modelling and analysis of intracellular metabolism. By assuming that the metabolic reaction network operates under steady state, the metabolic fluxes can be estimated from the measurements of extracellular metabolites using metabolic flux analysis (MFA) (when the estimation problem is over-determined) (Goudar et al., 2007) or using flux balance analysis (FBA) (when the estimation problem is under-determined) (Orth et al., 2010). In the FBA, one makes a further assumption that the cell organizes its metabolic flux distribution to optimize a particular objective (e.g. biomass production). While MFA and FBA have been routinely used to analyze metabolic network models, they are not yet commonly used to study protein glycosylation. A recent constraint-based modelling of protein glycosylation adopted a probabilistic approach using the Markov chain theory to explore feasible fluxes that could explain the observed glycan distribution, thereby avoiding the need to assign a cellular objective as in the FBA (Spahn et al., 2015). The method was developed specifically for predicting the protein glycosylation outcome from enzyme knock-out(s), and has been extended to glycoengineering by strain optimization (Spahn et al., 2017). While these works demonstrated the promise of CBM for glycoengineering application, the strategy was not designed for predicting the (dynamical) changes in the regulation of glycosylation fluxes due to alterations in the cell culture conditions and parameters (e.g., media compositions, pH, etc.).

In this work, we present a novel constraint-based modelling approach for analyzing protein glycosylation fluxes. The method, called Glycosylation Flux Analysis (GFA), generates predictions for the (intracellular) glycosylation reaction fluxes from the secretion rates of glycoforms. The GFA is based on the premise that the temporal variations in the intracellular glycosylation fluxes are the results of time-varying changes in the enzyme-specific factors (such as gene expression, enzyme activity and nucleotide sugar availability) and in the cell metabolism (especially the specific productivity). Finally, we applied the GFA to elucidate the dynamical changes of immunoglobulin G (IgG) glycosylation in fed-batch CHO cell cultures using two different media compositions (without and with manganese feed).

## 2. Materials and Methods

### 2.1 Cell culture conditions

CHO-S cells producing an IgG monoclonal antibody were cultivated in a 3 L bench scale bioreactor system (Sartorius Stedim, Germany). The resulting antibody bore two N-linked glycosylation sites at the Fc region. Cells were first expanded in shake bottles and the cell seeding density was set to 0.3×10^6^ cells/mL. A temperature shift from 36.5 °C to 33.0 °C and a pH shift from 7.1 to 6.9 were conducted on the fifth culture day. CO2 was used to control the pH and the dissolved oxygen set point was fixed to 50 % air saturation. Feeds were added on day 3, 5, 7 and 10 and consisted of a proprietary concentrated main feed with over 30 components, a highly alkaline amino acid solution and a glucose solution of 400 g/L. The main feed in the first experiment (process A) represented a control experiment, whereas the main feed in the second experiment (process B) had a different amino acid composition with manganese chloride addition. The final concentration of MnCl in process B was set to 2.4 μmol per liter culture volume.

### 2.2 Analytical methods for cell cultures and product quality analysis

Cell counts and cell viability were measured with a Vi-Cell analyzer (Bechman Coulter, Brea, CA), and metabolites were quantified with a Nova CRT (Nova Biomedical, Waltham, MA). A NOVA BioProfile pHOx analyzer (Nova Biomedical, Waltham, MA) was used to determine pH, pO2 and pCO2. The antibody concentration was determined on a Biacore C instrument (GE Healthcare, Waukesha, WI). Protein glycosylation of Phytips eluates (Phytips^®^, PhyNexus, San Jose, CA, USA) was analyzed by Ultra Performance Liquid Chromatography (UPLC)-2-amino-benzamide labelling technique. A 100 mm HILIC column was supplied by Waters Corporation, Milford, MA, USA. Samples at working day 3, 4, 5 and 6 were concentrated prior to analysis using 5 kDa MWCO vivaspins (Sartorius Stedim, Germany).

### 2.3 Data Pre-processing

The glycosylation flux analysis uses the secretion fluxes (rates) of protein glycoforms as inputs to make predictions on the intracellular glycosylation fluxes. For this purpose, the cell culture data of viable cell density (*X_v_*(*t*) [10^6^ cells/mL]), mAb concentration (*T*(*t*) [g/L]) and glycan fractions (*f_i_*(*t*) [%]) were pre-processed to produce the required secretion fluxes. First, the concentration of each glycoform was computed from the temporal data of mAb titer and the fraction of the glycoforms as follows:

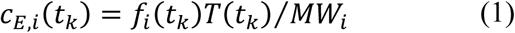
where *c_E,i_*(*t*_*k*_) denotes the extracellular concentration of the *i*-th glycoform at the *k*-th time point (*k* = 1, 2, …,*K*) and *MW_i_* denotes the molecular weight of the *i*-th glycoform. Since the contribution of glycans to the molecular weight of the protein was insignificant, the protein’s molecular weight was used for the calculations above (MW IgG = 150 kDa). The glycoform concentrations were smoothened using a sigmoidal Hill-type function:

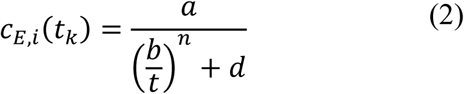
where the parameters *a, b, d* and *n* were determined by minimizing the sum of squares of the difference between the smoothing function and the concentration data. Finally, the time slopes of the glycoform concentration *dc_E,i_*(*t*)/*dt* were evaluated using the first-order derivative of the aforementioned smoothing function.

Meanwhile, the viable cell density data were smoothened using a logistic function, given by: (Goudar et al., 2005):

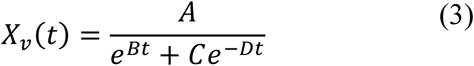
where the parameters *A, B, C* and *D* were obtained by minimizing the sum of squares of the difference between the logistic function above and *X_v_* data. Given *dc_E,i_*(*t*)*/dt* and *X_v_*(*t*) from the above data smoothening steps, the secretion fluxes of each glycoform *v_E,i_*(*t*) were computed according to:

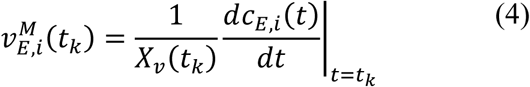

Note that in the above derivations, we assumed a constant volume of the cell culture. For the fed-batch experiments that we considered, such an assumption was reasonable since the cell culture volume varied only slightly with time (see SI Figure 1). When the volume of the cell culture varies strongly with time, the secretion flux calculation should take into account the volumetric changes, as follow:

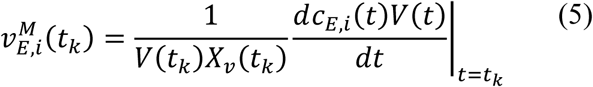
where *V*(*t*) is the cell culture volume at time *t*.

### 2.4 Glycosylation Flux Analysis

The glycosylation flux analysis (GFA) gives predictions for the intracellular fluxes of the glycosylation network (see Figure 1) under a pseudo steady state assumption based on a constraint-based modelling approach, as follow:

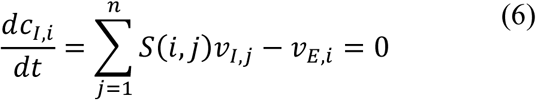
where *c*_*I,i*_ denotes the (internal) concentration of the *i*-th glycoform, *v*_*I,j*_ denotes the (internal) flux of the *j*-th glycosylation reaction, *v_E,i_* denotes the (external) secretion flux of the *i*-th glycoform, and *S*(*i*,*j*) is the number of molecules of the i-th glycoform that is consumed (when *S*(*i,j*) < 0) or produced (when *S*(*i,j*) > 0) by the *j*-th reaction. The pseudo steady state assumption is reasonable as the residence time of Golgi apparatuses between 20-40 minutes (Bibila and Flickinger, 1991; Hirschberg and Lippincott-Schwartz, 1999), is much shorter than the time scale of the cell culture dynamics (on the order of days). The relationship above can be written in a matrix-vector format as follows:

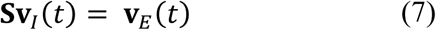
where **S** denotes the stoichiometric matrix of the network, and **v**_*I*_(*t*) and **v**_*E*_(*t*) denote the vectors of glycosylation and secretion fluxes of the protein glycoforms, respectively. The stoichiometric matrix is an *m* by *n* matrix where *m* is the number of protein glycoforms and *n* is the number of reactions in the network. Based on Eq. (7), the values of the glycosylation fluxes at each time point *t_k_* can be computed directly from the secretion fluxes of the glycoforms **v**_*E*_(*t*_*k*_) when the stoichiometric matrix **S** is invertible or has a full column rank. Given that the glycosylation enzymes can act on multiple substrates, the number of reaction exceeds the number of structures in the glycosylation network (i.e. *m* is smaller than *n*). In other words, the stoichiometric matrix is not invertible and the calculation of the internal glycosylation fluxes is an underdetermined problem.

**Figure 1.**
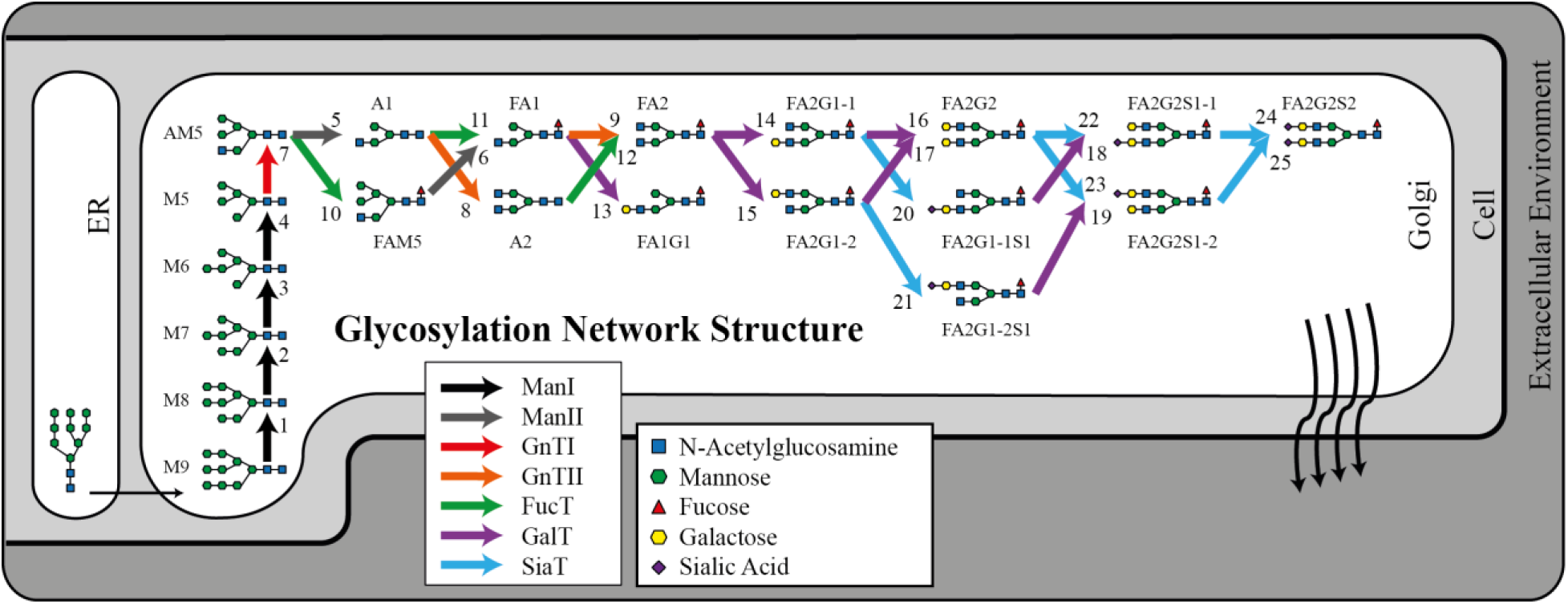
N-linked glycosylation network for immunoglobulin G in CHO-S. The glycosylation network model consists of 19 measured glycoforms (excluding M9) with 25 glycosylation reactions and represents the smallest network that includes all measured and intermediate glycoforms. The glycoform M9 is not required for the determination of intracellular glycosylation fluxes. The color of the arrows refers to the glycosylation enzymes which catalyze the corresponding flux.

In order to resolve the underdetermined problem above, we assume that the temporal changes in the glycosylation fluxes ***V***_*I*_ are the result of (i) local enzyme-specific alterations and (ii) global dynamical changes in the cell metabolism. Based on this assumption, we write the time evolution of internal glycosylation fluxes as follow:

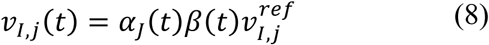
where *α*_*J*_(*t*) denotes the enzyme-specific (local) factor, *β*(*t*) denotes the cell specific (global) factor, and 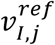 denotes the reference flux value for the *j*-th internal flux *v*_*I,j*_. The enzyme-specific factor *α*_*J*_(*t*) is shared among all reactions catalysed by the same enzyme *J*, while the cell factor *β*(*t*) applies to all reactions in the glycosylation network. The factors *α*_*J*_(*t*) and *β*(*t*) represent the time-dependent fold-amplification or -attenuation of the glycosylation fluxes, and hence their values can be normalized with respect to those at an arbitrary reference time point *t_ref_*. Without loss of generality, in this work we used the first measurement time point as the reference time, and thus assigned all *α*_*J*_(*t*_1_) and *β*(*t*_1_) to 1.

The factors *α*_*J*_(*t*) capture the dynamical alterations of glycosylation fluxes through local (enzyme-specific) factors such as enzyme expression, activity, inhibition, and nucleotide sugar availability. On the other hand, the variable *β*(*t*) describes the influence of the global cell metabolism on the glycosylation network, in particular the dynamical changes in the specific productivity of the cells. Therefore, we computed *β*(*t*) as the ratio between the specific productivity (*q_mAb_*) at time *t* and that at time *t_ref_*, as follows:

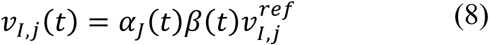

Given the relationships in Eqs. (8) and (9), the calculation of the internal glycosylation fluxes reduces to fitting *α*_*J*_(*t*) and 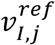 to the secretion flux values, according to:

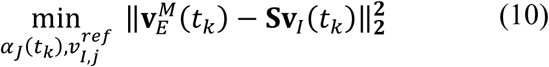

The schematic of the GFA workflow is depicted in Figure 2. The number of optimization variables above is *n*_*J*_(*K* − 1) + *n*, where *n*_*J*_ is the number of enzymes in the glycosylation network. As long as the total number of secretion flux values is larger than or equal to the number of unknowns above (i.e., *mK* ≥ *n*_*J*_(*K* − 1) + *n*), the GFA problem is over- or fully-determined, respectively. In this work, the optimization in the GFA was solved in MATLAB by using a global optimization method called the enhanced scatter search method from the MEIGO toolbox (Egea et al., 2014).

**Figure 2.**
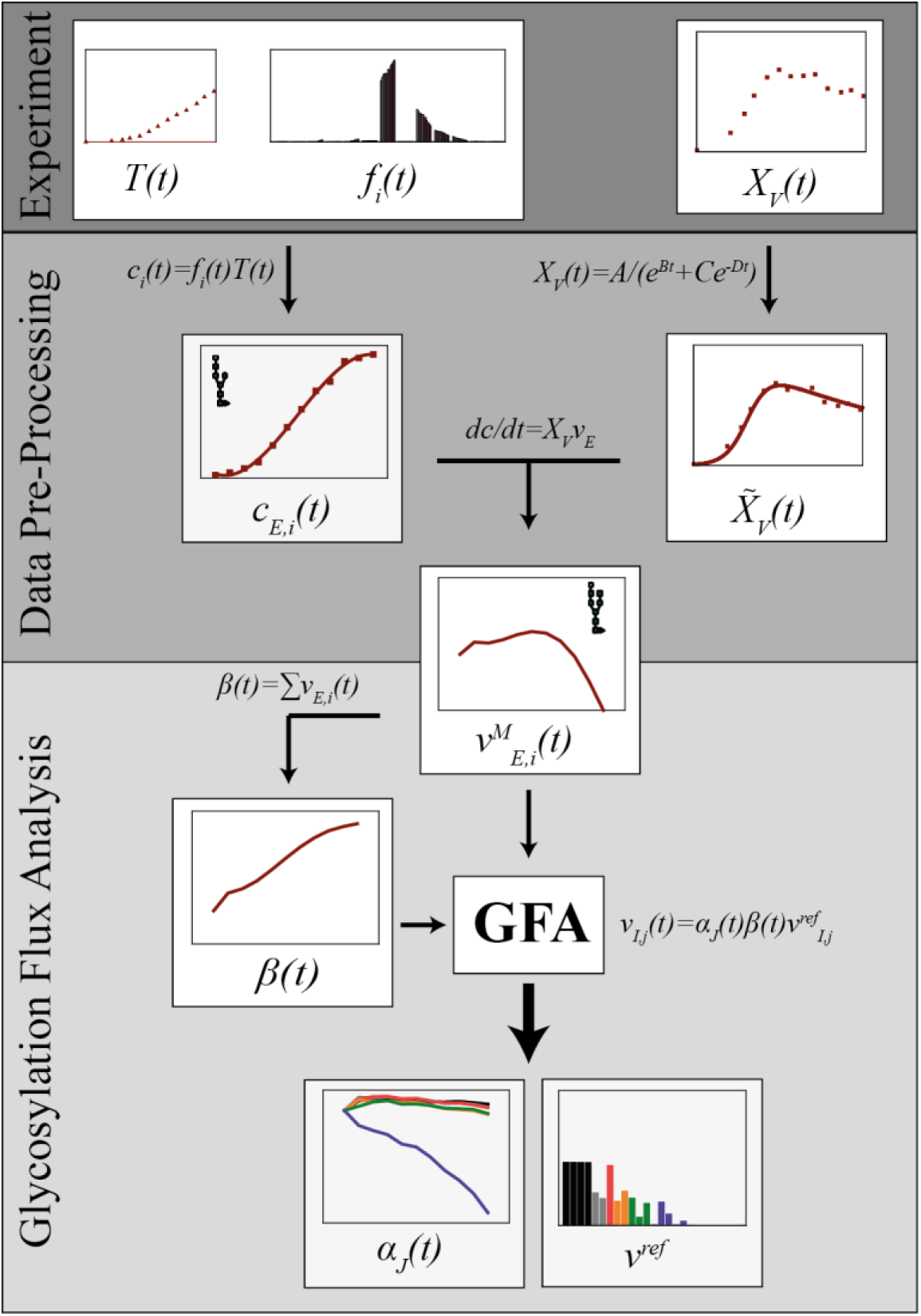
The GFA workflow. In the data pre-processing step, measurements of viable cell density (*X*_*v*_(*t*)), titer (*T*(*t*)) and the glycan fractions (*f*_*i*_(*t*)) are smoothened for the computation of the secretion fluxes of the glycoforms and the specific productivity of the cells *β*(*t*). The GFA predicts the intracellular fluxes by optimizing over the values of the enzyme specific factors *α*_*J*_(*t_k_*) and the reference flux values 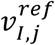.

For quantifying the uncertainty in the estimated *α*_*J*_(*t_k_*) and 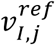, we generated 10 synthetic datasets for each process, consisting of glycoform concentrations and *X_v_* values at the same measurement time points, by contaminating the actual data with independent and identically distributed noise from a Gaussian distribution. We used the residuals from the data smoothening step to compute the variance of the noise for each variable. Finally, for each synthetic dataset, we carried out the GFA in the same manner as described above. In the application of the GFA to CHO fed-batch cultivation below, we reported the optimized *α*_*J*_(*t_k_*) and 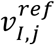 along with the standard deviation evaluated using the GFA results of the synthetic datasets.

## 3. Results

### 3.1 Cell culture data

Figure 3 shows the cell culture measurements of the viable cell density, IgG concentration and glycoform distribution (fractions) from process A and B. The logistic function smoothing for *X*_*v*_(*t*) appears as solid lines in the figure above. The cell density in process A reached a peak of ∼20×10^6^ cells/mL on day 7, while the peak cell density of process B was higher at ∼25×10^6^ cells/mL and occurred on day 8. While the viable cell density in process A was lower than that in process B, both processes had a similar final IgG concentration of ∼3 g/L. The logistic function could describe the overall temporal variation in the viable cell density data reasonably well (SI Table 1). A more detailed comparison between the two process datasets is provided in SI Figure 2.

**Figure 3.**
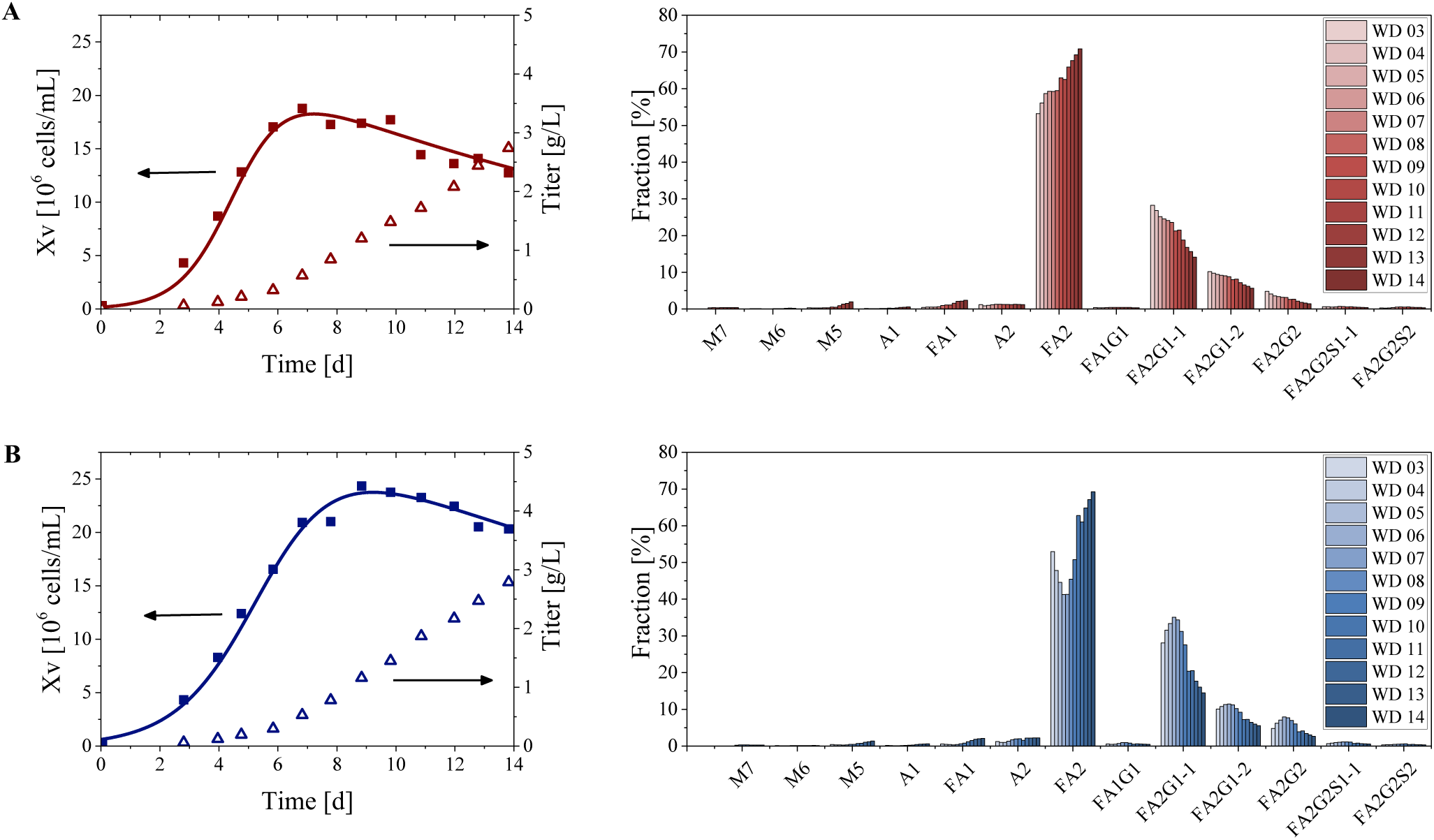
Cell culture measurements of the viable cell density (filled symbols), IgG titer (empty symbols) and glycoform distribution using (A, red) the standard culture media and (B, blue) the alternative media with Mn addition. The solid lines show the logistic function curve fitting for the viable cell density (see Methods).

The time evolution of the IgG glycan distribution in both processes showed an increase of less processed glycan fractions toward the end of the cell culture duration, in agreement with previous studies using the same cell line (Aghamohseni et al., 2014; Pacis et al., 2011). While such a trend was observed for the entire cell culture duration in process A, this trend was reversed in the first half of process B. More precisely, between day 3 and 6, we observed an increase in the fraction of galactosylated glycan structures. This observation agreed with a previous study investigating the effect of manganese addition using real-time glycosylation monitoring (Tharmalingam et al., 2015). Manganese is a cofactor of many Golgi resident enzymes, including N-acetylglucosaminyltransferases I/II and galactosyltransferases (Hendrickson and Imperiali, 1995).

### 3.2 GFA of CHO Fed-batch Cultivation

Figure 1 shows the glycosylation network used in the GFA of CHO-S cell culture. The network was derived from a previous publication (Villiger et al., 2016), in which undetectable and non-intermediate glycoforms were removed (Shah et al., 2008). In the GFA, we considered the glycosylation flux balances around all protein glycoforms downstream of M8 (including M8), which corresponded to *m* = 19 glycoforms and *n* = 25 reactions (see SI Text A for the glycosylation network model). For *K* = 11 time points, the number of secretion flux values (209) was larger than the number of unknowns (95), and hence the GFA was over-determined. A few of the glycoforms in the network (M8, AM5, FAM5, FA2G1-1S1, FA2G1-2S1, FA2G2S1-2) were undetected, and thus we set the corresponding secretion rates to 0. Figure 4 shows the concentration time profiles of the glycoforms and the corresponding Hill-type smoothing curves for the two process conditions.

**Figure 4.**
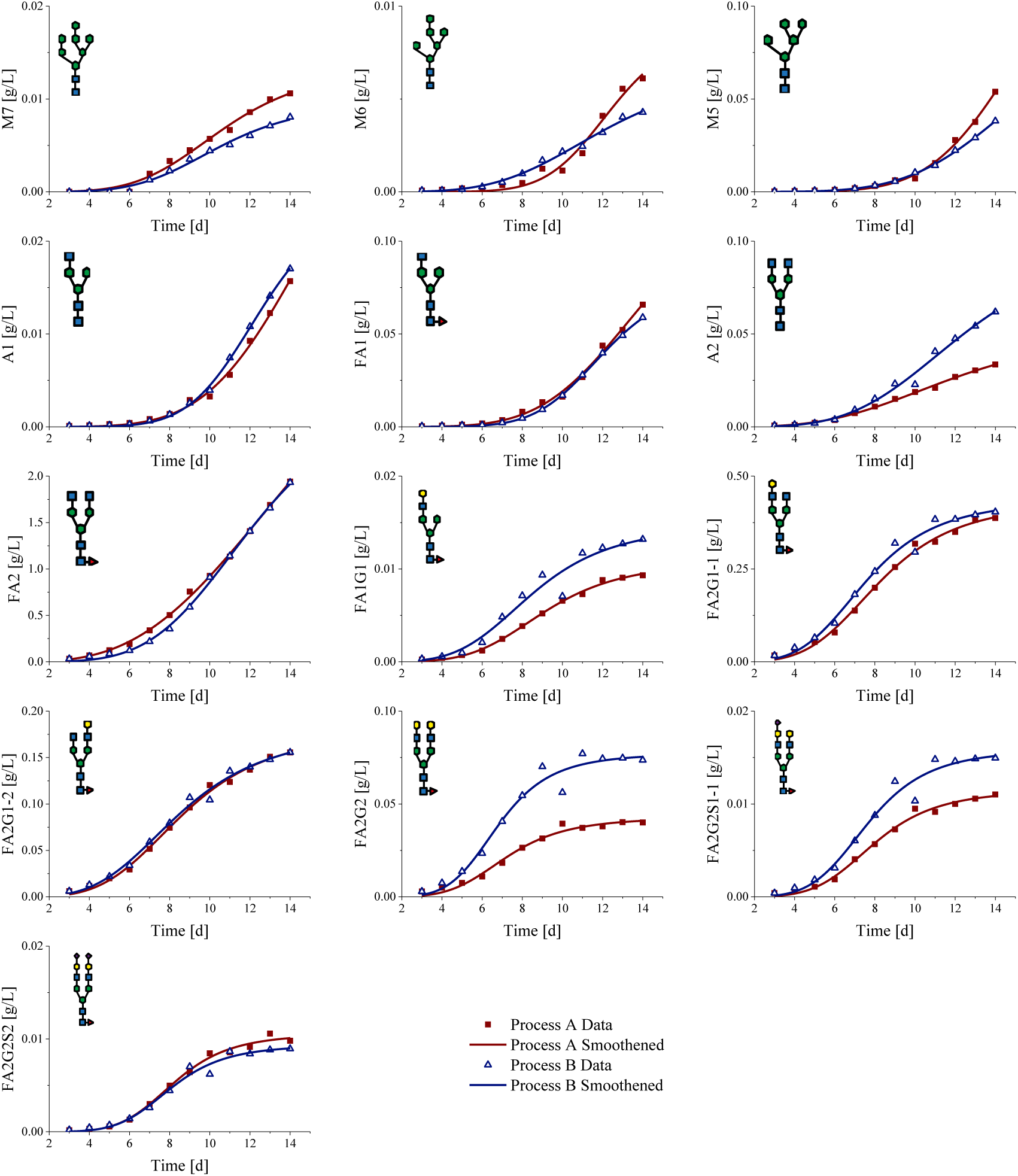
Calculated glycoform concentrations in process A (red, solid symbols) and process B (blue, empty symbols). The lines show the Hill-type smoothing function.

Figure 5 depicts the secretion fluxes of the glycoforms (in solid lines) after the data pre-processing. The secretion fluxes from the GFA are also drawn on the same figure, showing a good agreement with the smoothened flux values. As expected from the least square optimization in Eq. (10), the secretion fluxes of the main glycoforms with larger magnitudes, particularly FA2, were better reproduced by the GFA than others. While one could use a weighted least square optimization to obtain a better agreement with secretion fluxes of lower magnitudes, such a strategy would come at the cost of worse agreement for the fluxes producing the dominant glycan structures (results not shown).

**Figure 5.**
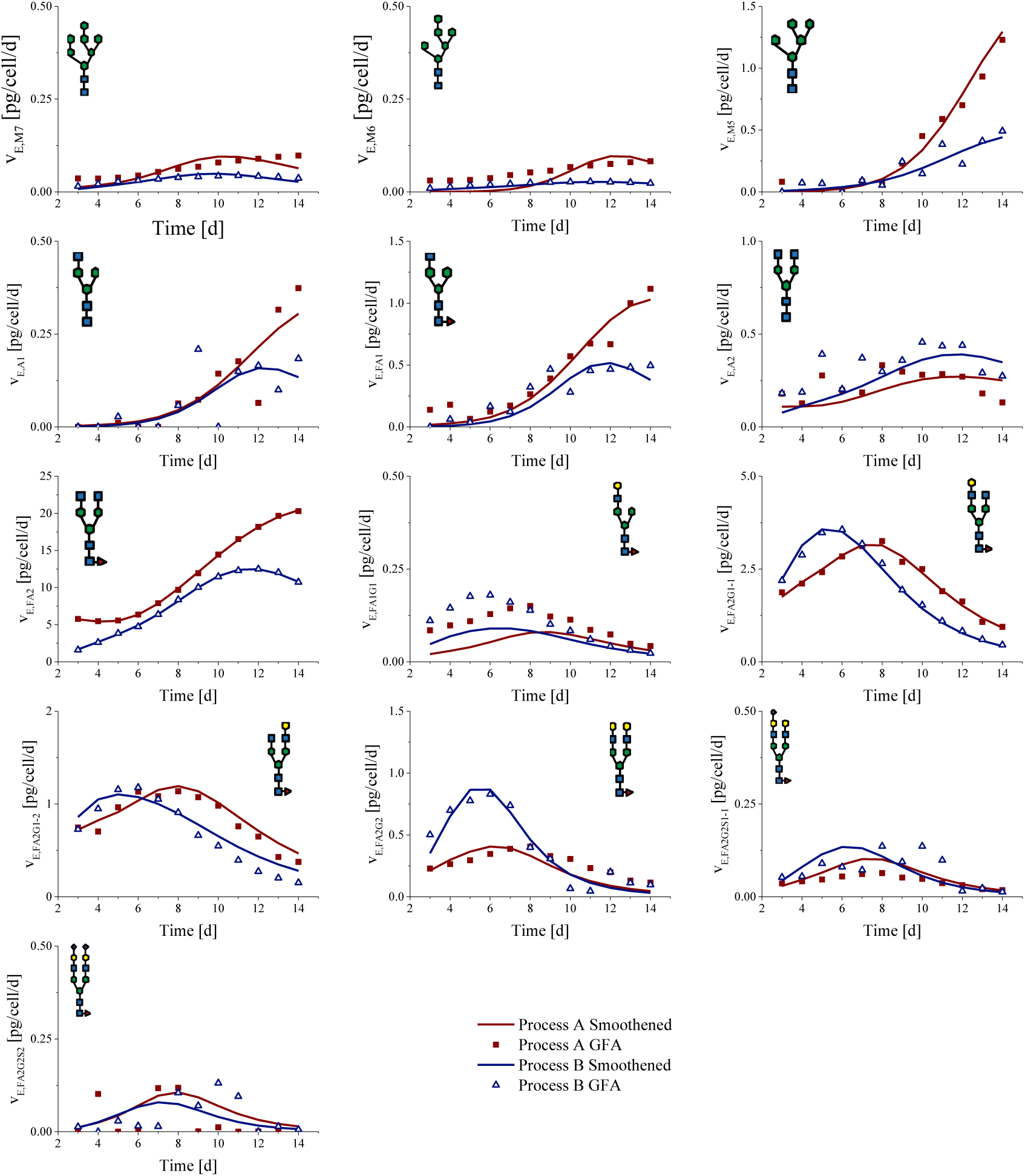
Fitting of glycoform secretion fluxes in GFA. The GFA gives accurate predictions of the secretory fluxes of glycoforms in process A (red, solid symbols) and process B (blue, empty symbols). The lines show the smoothened secretion fluxes in process A (red line) and process B (blue line).

Figure 6 describes the time profiles of the enzyme-specific factors *α*_*J*_(*t*), the cell specific productivity *β*(*t*), and the reference fluxes 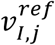 for process A and B. The cell specific productivity in the two processes generally increased with the cultivation time, with process A having a slightly higher fold-increase. The GFA predicted the enzyme-specific factor of galactosyltransferase GalT to decrease during much of the fed-batch cultivation period in both processes. The enzyme-specific factors of the other enzymes (ManI/II, GnTI/II, FucT) remained at an approximately constant level. A decrease in GalT activity is a frequently observed phenomena in industrial fed-batch processes of CHO-S culture (Gawlitzek et al., 2000), because of the adverse environmental conditions to which the cells are exposed (e.g., by-products, nutrient limitation and overfeeding, osmotic stress). As shown previously, the activity of GalT (and SiaT) for the cell line under investigation is much more sensitive to extracellular environment compared to ManI/II, GnTI/II and FucT (Villiger et al., 2016). In this regard, the accumulation of ammonia during the course of the cell culture (see SI Figure 2) could cause a pH increase in the Golgi, leading to a decrease in the GalT activity. The GFA of process A showed a slight increase in *α*_*GalT*_(*t*) between day 3 and 6, but such an increase was not statistically significant (H0: *α*_*GalT*_(*t*) = 1 has a *p* value larger than 0.05). Also, the drop in the GalT-specific factor explained the increase of FA2 and the decrease of galactosylated glycan structures towards the end of the two fed-batch cultivations. The flux values of reactions catalyzed by SiaT were not shown since these fluxes were predicted to be near zero (see SI Figure 3).

**Figure 6.**
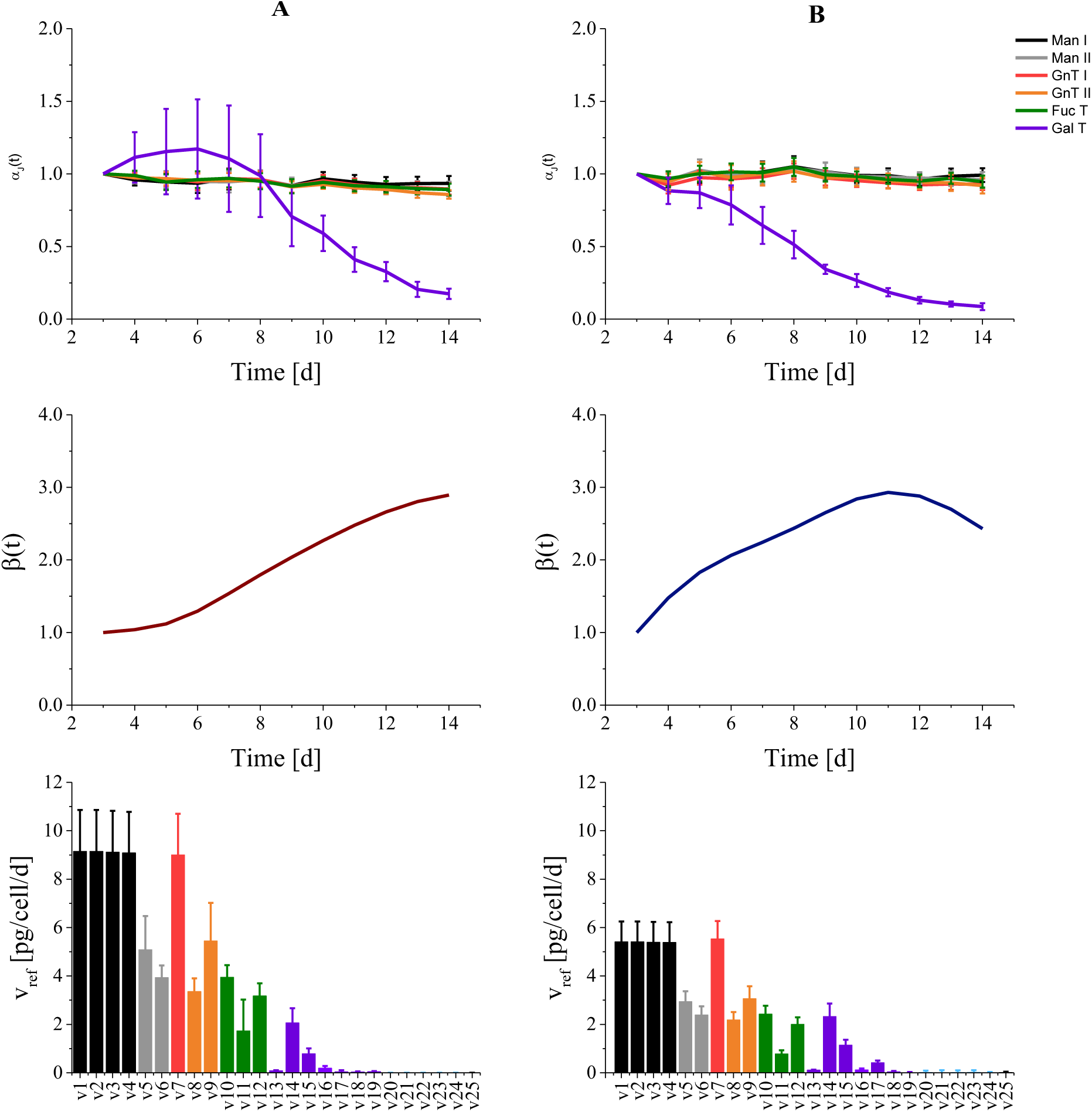
The GFA of process A and B: *α*_*J*_(*t*), *β*(*t*) and 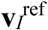. The error bars correspond to the standard deviation in the estimated variables (see Methods).

### 3.3 Effects of manganese

While processes A and B produced comparable glycoform distributions after day 10, the glycan fractions prior to this day, especially those of FA2 and the galactosylated structures, showed a completely different dynamic behavior. In order to elucidate the dynamic alterations in the IgG glycosylation caused by the change in the culture media composition and the addition of Mn, we computed the relative deviation in the viable cell density and in the (global) cell specific productivities (see Figure 7). The relative deviation between the two processes is computed as follows:

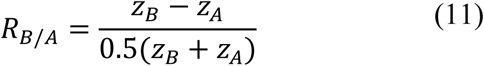
where *z_A_* and *z_B_* denote the variable of interest (i.e. the viable cell density and specific productivities) of process A and B, respectively. Figure 8 further shows the relative deviation of the normalized glycosylation throughput for each enzyme *J*, defined as

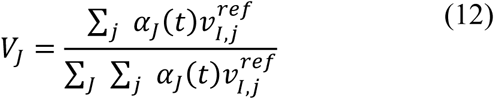

Note that the summation in the numerator above is done over all fluxes catalysed by the enzyme *J*. The normalized flux throughput of an enzyme reflects the activity of the glycosylation reactions carried out by a particular enzyme relative to the total activity of all glycosylation reactions in the network.

**Figure 7.**
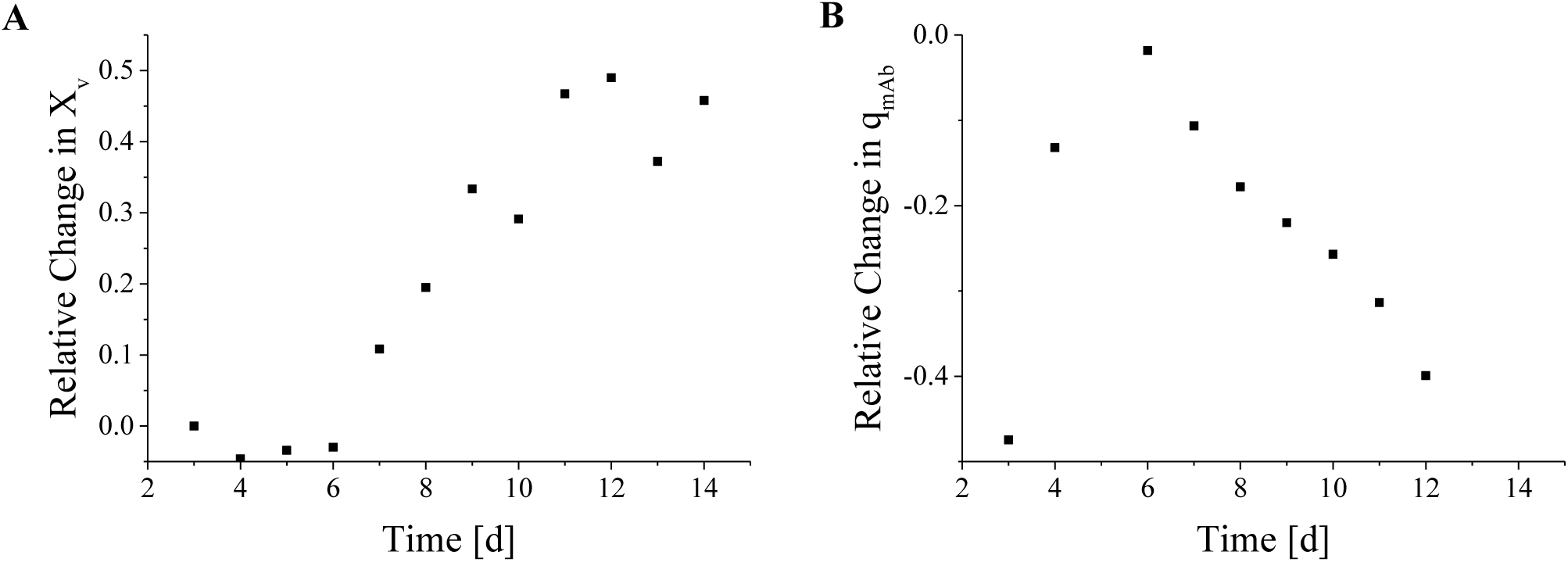
Relative deviations in (A) the viable cell density and (B) the cell specific productivity between process A and B. A positive (negative) value indicates that the variable is larger (smaller) in process B than in process A.

Figure 7 shows that the alternative culture media in process B led to a higher viable cell density but resulted in a lower specific productivity than process A. These observations suggested a trade-off between the cell growth and the antibody production upon changing the media composition. Except for fluxes catalysed by GalT, the normalized glycosylation throughputs in process B were roughly the same as those in process A for the entire cultivation period (see Figure 8). The normalized throughput of GalT showed a markedly more dynamic difference between the two processes. In particular, we observed a higher GalT normalized throughput in process B than in process A during the first half of the cell cultivation. This increase diminished with time and the relative deviation eventually dropped to roughly zero as with the other enzymes. Again, we did not compare the glycosylation fluxes catalysed by SiaT as the flux values were nearly zero (∼10^−3^ pg/cell/day).

**Figure 8.**
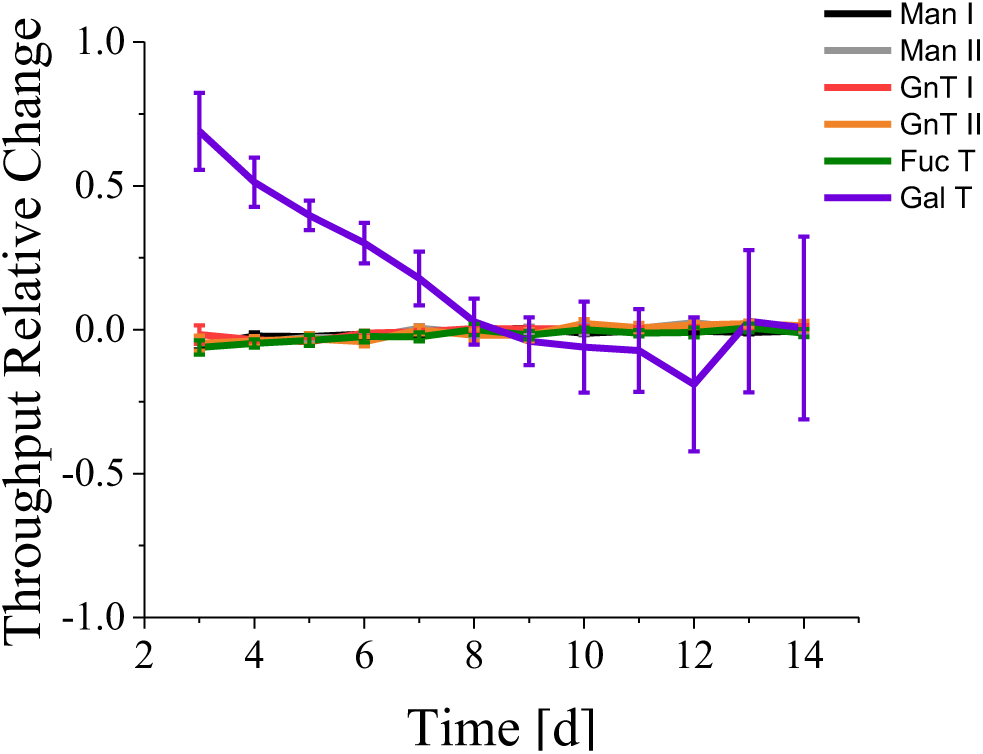
Relative deviations of the normalized flux throughput for each enzyme between process A and B. A positive (negative) value indicates that the variable is larger (smaller) in process B than in process A. The error bars correspond to the standard deviation in the estimated variables (see Methods).

## 4. Discussion

The combination of experimental and computational approaches, particularly using CBM, has brought a tremendous progress in metabolic engineering, for elucidating the regulation of cellular metabolism, for predicting the outcome of perturbations to the metabolic network (e.g. knock-outs), and for strain optimization (Bordbar et al., 2014). However, the CBM is not yet routinely applied to study the glycosylation network. Driven by the importance of understanding and controlling glycosylation of proteins, we created the glycosylation flux analysis by adapting the CBM approach for the glycosylation network. Based on a pseudo steady state assumption of the fluxes in the glycosylation network, the GFA generates dynamic glycosylation flux predictions using time-resolved measurements of cell culture data, particularly viable cell density, protein titer and glycan fractions. More specifically, the GFA captures the dynamic changes in the glycosylation fluxes using two pre-multiplicative factors; *α*_*J*_(*t*) that accounts for the (local) changes that occur in an enzyme-specific manner, and *β*(*t*) that accounts for the (global) alterations in cell metabolism that affects the cell specific productivity. Besides reducing the degrees of freedom in the flux prediction problem (due to the small number of enzymes in the glycosylation network), the use of pre-multiplicative factors above leads to the delineation of local and global effects of the process conditions on the protein glycosylation. In contrast to the more detailed differential equation based analyses of the glycosylation networks (Jedrzejewski et al., 2014; Jimenez del Val et al., 2011; Krambeck and Betenbaugh, 2005), the GFA, like the MFA, represents a parameter-free approach and requires only the stoichiometric matrix information. However, since the GFA involves solving the inverse problem – estimating intracellular fluxes given dynamic cell culture data, this method cannot directly be used for predicting the effects of changing process parameters or genetic alterations on *α*_*J*_(*t*), *β*(*t*) and 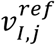 *a priori*.

There exist several implications from the formulation of intracellular glycosylation flux *V_I,j_* that depends on three variables: *α*_*J*_(*t*), *β*(*t*) and 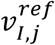. Starting with *β*(*t*), for fixed values of *α*_*J*_(*t*), a fold-increase (-decrease) in *β*(*t*) means an equal fold-increase (-decrease) among all of the glycosylation and secretion fluxes in the network (see SI Text B). Therefore, *β*(*t*) captures the global change in the glycosylation network, but such a change will not affect the glycan distribution in the monoclonal antibody. Meanwhile, for fixed values of *β*(*t*) and 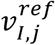, a variation in the enzyme-specific factors *α*_*J*_(*t*) will lead to a reorganization of the glycosylation fluxes and thus changes in the glycan distribution. The reorganization of the internal fluxes is restricted by constraints enforced by the flux balance equation in Eq. (7) (see SI Text B).Because all intracellular glycosylation fluxes associated with a particular enzyme *J* scale equally by *α*_*J*_(*t*), the ratios among these fluxes thus remain constant with variations in *α*_*J*_(*t*). Such ratios are given by the relative magnitudes of the corresponding 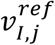(see Figure 6). Finally, since *α*_*J*_(*t*) and *β*(*t*) are set to 1 at the reference time point *t*_*ref*_, 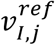 are the predictions for the intracellular glycosylation fluxes at this time point, i.e. 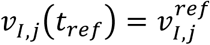.

We applied the GFA to elucidate the glycosylation fluxes in two fed-batch cultivations of CHO cells producing IgG using different cell culture media. Our analyses showed that the temporal variation in the glycan distribution mainly resulted from the decrease in the galactosylation activity with time, relative to the other glycosylation reactions. We attributed this drop to the sensitivity of GalT activity to environmental changes, particularly the accumulation of ammonia in the cell culture. While the time profiles of the viable cell density and the specific productivity of IgG were similar between the two fed-batch processes, the changes in the glyoprofiles during the first half of the cultivations followed markedly different dynamics. The GFA showed that between day 3 and 6, the addition of Mn, a co-factor of GalT (Ramakrishnan et al., 2004). in process B increased the normalized galactosylation throughput. However, this increase largely disappeared by day 10. While Mn has also been reported to be a co-factor for GnTI/II (Nishikawast et al., 1988), under these cultivation conditions, the normalized glucosamination throughput in process B was roughly equal in that in process A. This observation suggested that the enzyme activity of GnTI/II might not be the rate-limiting factor of the IgG glucosamination in the two fed-batch cultivations.

## 5. Conclusion

In this work, we adapted a constraint-based modeling approach to the analysis of protein glycosylation and developed the glycosylation flux analysis method. The GFA uses extracellular data of glycoform secretion fluxes and a (pseudo) steady state model of the glycosylation network to determine the intracellular glycosylation fluxes. In this regard, the GFA separates the influence of local enzyme-specific factors and global cell metabolism on each glycosylation flux. The ability to decouple the influences of local and global factors is important in elucidating the regulation of protein glycosylation and especially in efforts to control the outcome of the glycosylation process. The application of the GFA to two fed-batch cultivations of CHO cells for IgG production demonstrated the dynamic information that one could obtain from this analysis, specifically on the key glycosylation enzyme(s) that governed the resulting glycoform distribution. As more research efforts are invested in understanding protein glycosylation, producing a treasure trove of cell culture and glycoprofiling data, a network-based analysis method, like the GFA, will play an enabling role in generating insights into the complex intracellular regulation and control of the protein glycosylation process.

## Acknowledgments

The authors would like to thank Daniel Karst, Ernesto Scibona and Prof. Massimo Morbidelli as well as Robert Gauss and Prof. Markus Aebi for fruitful discussions. MS was partially supported by the Specific University Research grant number 20/2015. SH was supported by funding from ETH Zurich.

## Symbols

*α*: Enzyme specific factor
*β*: Specific productivity
*c*: Concentration
*c_E_*: Extracellular concentration
*f*: Fraction
*K*: Number of time points
*m*: Number of glycoforms
*MW*: Molecular weight
*n*: Number of reactions
*S*: Stoichiometric matrix
*T*: Titer
t: Time
*V*: Volume
*V_E_*: Secretion flux
*V_I_*: Intracellular flux
*V^ref^I*: Reference fluxX
*X_v_*: Viable cell density

## Abbreviations

CBM: Constraint-based modelling
CHO: Chinese hamster ovarian
ER: Endoplasmic reticulum
FBA: Flux balance analysis
GFA: Glycosylation flux analysis
IgG: Immunoglobulin G
MFA: Metabolic flux analysis
PAT: Process analytical technology
QbD: Quality by design
UPLC: Ultra performance liquid chromatography

## Glycosylation nomenclature

Man: Mannosidase
GnT: N-Acetylglucosaminyltransferase
FucT: Fucosyltransferase
GalT: Galactosyltransferase
SiaT: Sialyltransferase
M9: Man_9_GlcNAc_2_
M8: Man_8_GlcNAc_2_
M7: Man_7_GlcNAc_2_
M6: Man_6_GlcNAc_2_
M5: Man_5_GlcNAc_2_
A1: GlcNAcMan_3_GlcNAc_2_
A2: GlcNAc_2_Man_3_GlcNAc_2_
FA1: GlcNAcMan_3_GlcNAc_2_Fuc
FA2: GlcNAc_2_Man_3_GlcNAc_2_Fuc
FA1G1: GalGlcNAcMan_3_GlcNAc_2_Fuc
FA2G1-1: α(1-6)GalGlcNAc_2_Man_3_GlcNAc_2_Fuc
FA2G2S1-2: α(1 -3)GalGlcNAc_2_Man_3_GlcNAc_2_Fuc
FA2G2: Gal_2_GlcNAc_2_Man_3_GlcNAc_2_Fuc
FA2G2S1-1: α(1 -6)SiaGal_2_GlcNAc_2_Man_3_GlcNAc_2_Fuc
FA2G2S1-2: α(1-3)SiaGal_2_GlcNAc_2_Man_3_GlcNAc_2_Fuc

## References

Aebi, M., 2013. N-linked protein glycosylation in the ER. Biochim. Biophys. Acta - Mol. Cell Res. 1833, 2430–2437. doi:10.1016/j.bbamcr.2013.04.001

Aebi, M., Hennet, T., 2001. Congenital disorders of glycosylation: genetic model systems lead the way. Trends Cell Biol. 11, 136–141. doi:10.1016/S0962-8924(01)01925-0

Aghamohseni, H., Ohadi, K., Spearman, M., Krahn, N., Moo-Young, M., Scharer, J.M., Butler, M., Budman, H.M., 2014. Effects of nutrient levels and average culture pH on the glycosylation pattern of camelid-humanized monoclonal antibody. J. Biotechnol. 186, 98–109. doi:10.1016/j.jbiotec.2014.05.024

Arosio, P., Rima, S., Morbidelli, M., 2013. Aggregation mechanism of an IgG2 and two IgG1 monoclonal antibodies at low pH: from oligomers to larger aggregates. Pharm. Res. 30, 641–654. doi:10.1007/s11095-012-0885-3

Bibila, T.A., Flickinger, M.C., 1991. A model of interorganelle monoclonal antibody transport and secretion in mouse hybridoma cells. Biotechnol. Bioeng. 38, 767–780. doi:10.1002/bit.260380711

Bordbar, A., Monk, J.M., King, Z.A., Palsson, B.O., 2014. Constraint-based models predict metabolic and associated cellular functions. Nat. Rev. Genet. 15, 107–20. doi:10.1038/nrg3643

Egea, J.A., Henriques, D., Cokelaer, T., Villaverde, A.F., MacNamara, A., Danciu, D.-P., Banga, J.R., Saez-Rodriguez, J., 2014. MEIGO: an open-source software suite based on metaheuristics for global optimization in systems biology and bioinformatics. BMC Bioinformatics 15, 136. doi:10.1186/1471-2105-15-136

FDA, 2004. Guidance for Industry. PAT — A Framework for Innovative Pharmaceutical DEvelopment, Mannufacturing, and Quality Assurance 19.

Gawlitzek, M., Ryll, T., Lofgren, J., Sliwkowski, M.B., 2000. Ammonium alters N-glycan structures of recombinant TNFR-IgG: degradative versus biosynthetic mechanisms. Biotechnol. Bioeng. 68, 637–646. doi:10.1002/(SICI)1097-0290(20000620)68:6<637::AID-BIT6>3.0.CO;2-C

Goh, J.S.Y., Liu, Y., Liu, H., Chan, K.F., Wan, C., Teo, G., Zhou, X., Xie, F., Zhang, P., Zhang, Y., Song, Z., 2014. Highly sialylated recombinant human erythropoietin production in large-scale perfusion bioreactor utilizing CHO-gmt4 (JW152) with restored GnT I function. Biotechnol. J. 9, 100–109. doi:10.1002/biot.201300301

Goudar, C.T., Biener, R., Piret, J.M., Konstantinov, K.B., 2007. Metabolic flux estimation in mammalian cell cultures, in: Methods in Biotechnology. Springer, pp. 301–317. doi:10.1002/btpr.284

Goudar, C.T., Joeris, K., Konstantinov, K.B., Piret, J.M., 2005. Logistic equations effectively model Mammalian cell batch and fed-batch kinetics by logically constraining the fit. Biotechnol. Prog. 21, 1109–18. doi:10.1021/bp050018j

Harding, F.A., Stickler, M.M., Razo, J., DuBridge, R.B., 2010. The immunogenicity of humanized and fully human antibodies: Residual immunogenicity resides in the CDR regions. MAbs 2, 256–265. doi:10.4161/mabs.2.3.11641

Hendrickson, T.L., Imperiali, B., 1995. Metal ion dependence of oligosaccharyl transferase: implications for catalysis. Biochemistry 34, 9444–9450. doi:10.1021/bi00029a020

Hirschberg, K., Lippincott-Schwartz, J., 1999. Secretory pathway kinetics and in vivo analysis of protein traffic from the Golgi complex to the cell surface. Faseb J 13 Suppl 2, S251–S256.

Hossler, P., Khattak, S.F., Li, Z.J., 2009. Optimal and consistent protein glycosylation in mammalian cell culture. Glycobiology 19, 936–949. doi:10.1093/glycob/cwp079

Ivarsson, M., Villiger, T.K., Morbidelli, M., Soos, M., 2014. Evaluating the impact of cell culture process parameters on monoclonal antibody N-glycosylation. J. Biotechnol. 188C, 88–96. doi:10.1016/j.jbiotec.2014.08.026

Jedrzejewski, P.M., del Val, I.J., Constantinou, A., Dell, A., Haslam, S.M., Polizzi, K.M., Kontoravdi, C., 2014. Towards controlling the glycoform: A model framework linking extracellular metabolites to antibody glycosylation. Int. J. Mol. Sci. 15, 4492–4522. doi:10.3390/ijms15034492

Jimenez del Val, I., Nagy, J.M., Kontoravdi, C., 2011. A dynamic mathematical model for monoclonal antibody N-linked glycosylation and nucleotide sugar donor transport within a maturing Golgi apparatus. Biotechnol. Prog. 27, 1730–43. doi:10.1002/btpr.688

Krambeck, F.J., Betenbaugh, M.J., 2005. A mathematical model of N-linked glycosylation. Biotechnol. Bioeng. 92, 711–728. doi:10.1002/bit.20645

Li, F., Vijayasankaran, N., Shen, A., Kiss, R., Amanullah, A., 2010. Cell culture processes for monoclonal antibody production. MAbs 2, 466–479. doi:10.4161/mabs.2.5.12720

Liu, G., Puri, A., Neelamegham, S., 2013. Glycosylation Network Analysis Toolbox: a MATLAB-based environment for systems glycobiology. Bioinformatics 29, 404–406. doi:10.1093/bioinformatics/bts703

Nishikawast, Y., Peggs, W., Paulsenll, H., Schachters, H., 1988. Control of glycoprotein synthesis. Purification and characterization of rabbit liver UDP-N-acetylglucosamine: alpha-3-D-mannoside beta-1,2-N-acetylglucosaminyltransferase I. J. Biol. Chem. 263, 8270–8281.

Orth, J.D., Thiele, I., Palsson, B.Ø., 2010. What is flux balance analysis? Nat. Biotechnol. 28, 245–248. doi:10.1038/nbt.1614.What

Pacis, E., Yu, M., Autsen, J., Bayer, R., Li, F., 2011. Effects of cell culture conditions on antibody N-linked glycosylation-what affects high mannose 5 glycoform. Biotechnol. Bioeng. 108, 2348–2358. doi:10.1002/bit.23200

Ramakrishnan, B., Boeggeman, E., Ramasamy, V., Qasba, P.K., 2004. Structure and catalytic cycle of beta-1,4-galactosyltransferase. Curr. Opin. Struct. Biol. 14, 593–600. doi:10.1016/j.sbi.2004.09.006

Rathore, A.S., 2009. Roadmap for implementation of quality by design (QbD) for biotechnology products. Trends Biotechnol. 27, 546–553. doi:10.1016/j.tibtech.2009.06.006

Sha, S., Agarabi, C., Brorson, K., Lee, D.-Y., Yoon, S., 2016. N-Glycosylation Design and Control of Therapeutic Monoclonal Antibodies. Trends Biotechnol. 34, 835–846. doi:10.1016/j.tibtech.2016.02.013

Shah, N., Kuntz, D.A., Rose, D.R., 2008. Golgi alpha-mannosidase II cleaves two sugars sequentially in the same catalytic site. Proc. Natl. Acad. Sci. U. S. A. 105, 9570–5. doi:10.1073/pnas.0802206105

Spahn, P.N., Hansen, A.H., Henning, G., Arnsdorf, J., Kildegaard, H.F., Lewis, N.E., Arnsdorf, J., Kildegaard, H.F., Lewis, N.E., Markov, A., 2015. A Markov chain model for N-linked protein glycosylation – towards a low-parameter tool for model-driven. Metab. Eng. 1–15. doi:10.1016/j.ymben.2015.10.007

Spahn, P.N., Hansen, A.H., Kol, S., Voldborg, B., Lewis, N.E., 2017. Predictive glycoengineering of biosimilars using a Markov chain glycosylation model. Biotechnol. J. 12, 1–8.doi:10.1002/biot.201600489

St Amand, M.M., Tran, K., Radhakrishnan, D., Robinson, A.S., Ogunnaike, B. a, 2014. Controllability analysis of protein glycosylation in CHO cells. PLoS One 9, e87973. doi:10.1371/journal.pone.0087973

Tharmalingam, T., Wu, C.-H., Callahan, S., T Goudar, C., 2015. A framework for real-time glycosylation monitoring (RT-GM) in mammalian cell culture. Biotechnol. Bioeng. 112, 1146–1154. doi:10.1002/bit.25520

Umaña, P., Bailey, J.E., 1997. A Mathematical Model of N-Linked Glycoform Biosynthesis. Biotechnol. Bioeng. 55, 890–908. doi:10.1002/(SICI)1097-0290(19970920)55:6<890::AID-BIT7>3.0.CO;2-B

Umaña, P., Jean-Mairet, J., Moudry, R., Amstutz, H., Bailey, J.E., 1999. Engineered glycoforms of an antineuroblastoma IgG1 with optimized antibody-dependent cellular cytotoxic activity. Nat. Biotechnol. 17, 176–80. doi:10.1038/6179

Villiger, T.K., Scibona, E., Stettler, M., Broly, H., Morbidelli, M., Soos, M., 2016. Controlling the time evolution of mAb N-linked glycosylation - Part II: Model-based predictions.Biotechnol. Prog. doi:10.1002/btpr.2315

Zhang, K., Kaufman, R.J., 2006. Protein folding in the endoplasmic reticulum and the unfolded protein response., in: Handbook of Experimental Pharmacology. pp. 69–91.

